# The Role of HERG channel in the Secretion of Glucagon-Like Peptide-1 (GLP-1) from Murine Intestinal L-Cells

**DOI:** 10.1101/2023.02.28.530535

**Authors:** Chang Liu, Ying-Chao Yuan, Rong-Rong Xie, Hao Wang, Jin-Kui Yang

**Author notes:** Correspondence: Hao Wang or Jin-Kui Yang. These authors contributed equally: Chang Liu and Ying-Chao Yuan.

## Abstract

HERG ion channel is a member of the Voltage-gated potassium (Kv) channels. A reduction in HERG function reduces potassium efflux during repolarization. Previous research has shown that patients with long QT syndrome due to HERG mutations have increased secretion of the hormone glucagon-like peptide-1 (GLP-1). However, the role of HERG in GLP-1 secretion remains uncertain. Here we report that HERG is expressed in GLP-1-producing L-cells in rodent intestinal epithelium. In a mouse L-cell model (GLUTag cell line), downregulation of HERG significantly prolonged action potential duration, increased intracellular calcium concentration, and stimulated GLP-1 secretion after exposure to nutrients. These findings suggest that HERG in the intestine plays a direct role in GLP-1 secretion and may be a potential target for diabetes treatment.

## Background

The human ether-a-go-go-related gene (HERG) encodes the HERG ion channel, a member of the voltage-dependent potassium channel (Kv) family. Mutations in HERG lead to reduced potassium efflux during repolarization and prolonged QT interval, resulting in polymorphic ventricular tachycardia, cardiac syncope, and sudden death (1, 2). This condition is known as Long QT Syndrome (LQTS)(3). The HERG ion channels which are members of the larger Kv channel family(4), are expressed in the human pancreas and have been shown to negatively regulate insulin secretion and positively regulate glucagon secretion (5). Previous studies have demonstrated that patients with LQTS due to loss-of-function mutations in HERG showed elevated GLP-1 secretion after an oral glucose tolerance test (OGTT) (2). However, the role of HERG in L-cells remains unexplored.

Repolarizing potassium channels play a crucial role in hormone secretion in endocrine cells (5). In pancreatic beta cells, the conversion of glucose into ATP closes the KATP channels and depolarizes the cell membrane (6, 7). This results in an influx of calcium, which stimulates insulin exocytosis (5, 8). The depolarization of the membrane also triggers the opening of repolarizing (voltage-gated) Kv channels, allowing the action potential to repolarize. Our previous research has shown that insulin secretion in pancreatic islet beta cells is regulated by the interplay of both depolarized KATP channels and repolarized Kv channels during glucose stimulation (9). Inhibiting repolarization of Kv channels has been shown to enhance insulin secretion in beta cells (10, 11).

Glucagon-like peptide-1 (GLP-1) is a hormone released by intestinal L-cells that stimulates insulin release in response to food intake. The glucose sensing mechanism in L-cells is similar to that found in pancreatic cells (12). Glucose uptake in intestinal epithelial cells is facilitated by brush-edge sodium-glucose co-transporters, allowing glucose to cross the apical membrane into the intestinal L cells. Intracellular ATP generation then closes ATP-dependent potassium ion channels, depolarizing the cell membrane and allowing calcium influx, which triggers GLP-1 release (13, 14). Due to the recent success of GLP-1 mimetics and inhibitors of GLP-1 degradation in treating type 2 diabetes (15), there is increasing interest in targeting L-cells to promote GLP-1 secretion (16, 17). Understanding the sensory and secretory pathways of L-cells is crucial for the success of this strategy.

In this study, we examined the role of HERG in L-cells from the murine GLUTag cell line. Our results showed that HERG channels are highly expressed in GLUTag cells. When HERG was knocked down, there was a significant reduction in Kv currents and prolongation of the action potential duration. Furthermore, reducing HERG expression increased the intracellular calcium concentration ([Ca2+]i) and enhanced GLP-1 secretion from GLUTag cells following nutrient stimulation. These findings reveal that the HERG channel plays a regulatory role in GLP-1 secretion in murine L-cells.

## Methods

### Animal care and cell culture

C57BL/6J mice were obtained from Jiangsu Gempharmatech Laboratories (Jiangsu, Nanjing, China). These mice were kept in a controlled environment with a consistent temperature and humidity, a 12-hour light-dark cycle, and were fed a standard diet. Only male mice were used in the experiments and all animal testing was performed in compliance with the regulations set by the Animal Care and Experimentation Committee at Capital Medical University. The GLUTag cell line, which has been validated as a model for GLP-1 secretion by intestinal L-cells, was used in our experiments (12, 18). GLUTag cells were cultured in high-glucose Dulbecco’s modified Eagle medium (DMEM) with a glucose concentration of 25 mM and supplemented with 10% fetal bovine serum (FBS). The cells were grown in a humidified incubator with 95% air and 5% CO2 at a temperature of 37°C.

### Immunoblotting analysis

For protein extraction, tissues and cells were lysed in a buffer consisting of 20 mM Tris-HCl pH 7.5, 150 mM NaCl, 1 mM MgCl2, 1% Triton X-100, 1 mM PMSF, and a complete protease inhibitor cocktail (Roche). The resulting supernatants, containing the proteins, were collected after centrifugation at 14,000 rpm for 10 minutes at 4°C. Western blot analysis was then performed according to a previously described protocol (19). The immunoreactive signals were detected using ECL Prime (GE Healthcare Biosciences) and an ImageQuant 500 chemiluminescence detection system (GE Healthcare Bioscience).

### Immunofluorescence staining

To study the cellular localization of proteins, immunofluorescence staining was performed on frozen sections from C57BL/6N mice. The sections were fixed with 3% paraformaldehyde in phosphate-buffered saline (PBS) for 30 minutes and permeabilized with 0.1% Triton X-100 in PBS for an additional 30 minutes. The cells were then blocked with PBS containing 1% bovine serum albumin (BSA) for 15 minutes. Primary antibody incubation was performed overnight, followed by 60 minutes of incubation with Alexa Fluor 488-or 568-conjugated secondary antibody (Invitrogen, at a dilution of 1:500). The images were captured using a 3D Histech Digital Pathology System and CaseViewer software.

### RNA isolation and gene expression analysis

RNA extraction was performed using Trizol Reagent (Invitrogen) as per the manufacturer’s protocol. 1 μg of total RNA was reverse transcribed using an oligo-(dT)12-18 primer and Superscript III (Invitrogen). Quantitative PCR gene expression analysis was conducted using a LightCycler 480 Real-Time PCR System (Roche, Basel, Switzerland) with SYBR Green I Master Mix reagent (Roche) and the primers listed in Table 1. All reactions were performed in triplicate and the relative mRNA expression levels were calculated and normalized against the expression of Rplp0/36B4 mRNA.

**Table 1.**
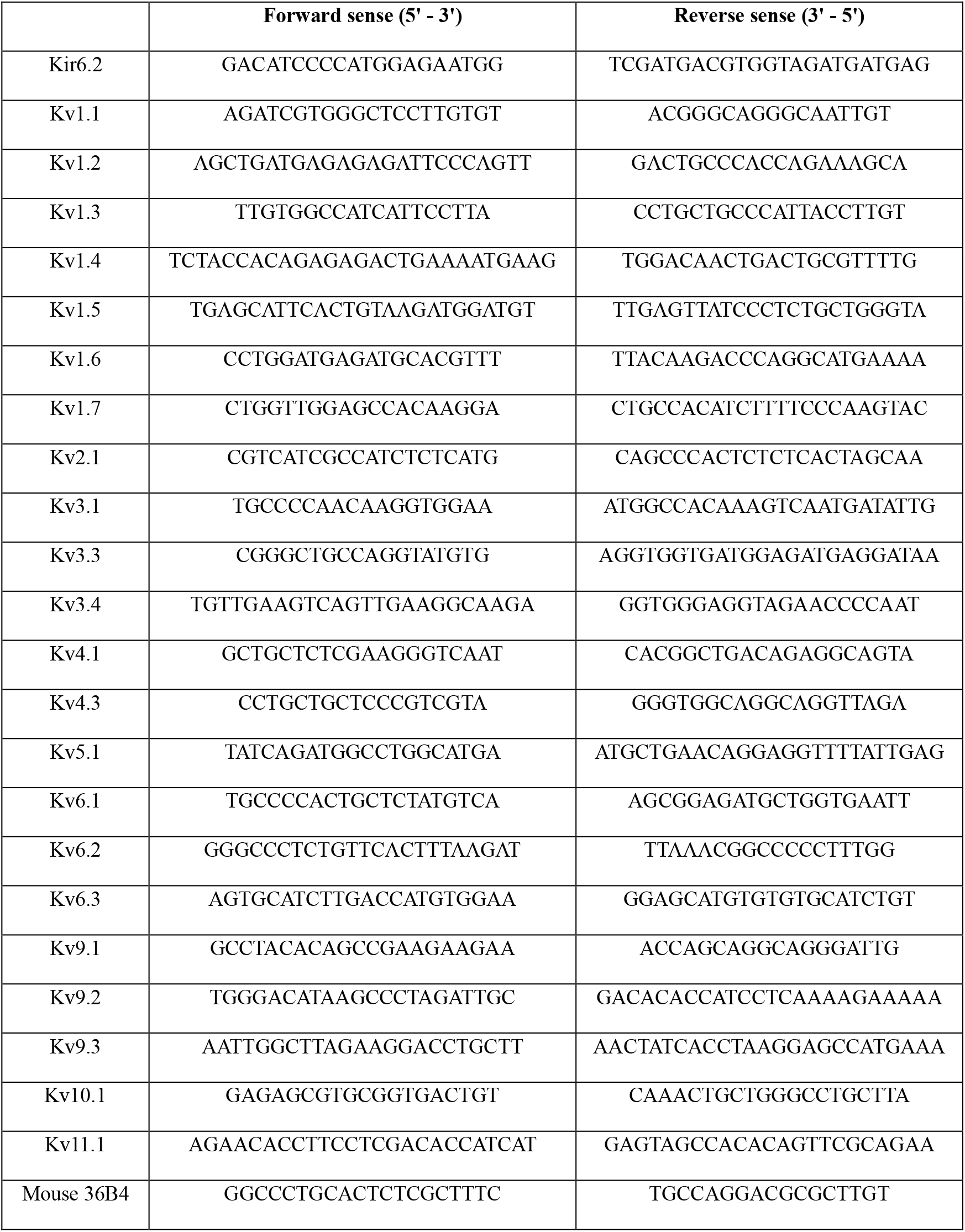
qPCR primers of mouse Kv channels and 36B4.

### siRNA transfection

GLUTag cells were seeded in 24-well plates and when they reached 80% confluency, they were transfected with 100 nM scramble siRNA or siRNA against mouse kcnh2 (sc-42498, Santa Cruz) using Lipofectamine RNAiMAX (Invitrogen) as per the manufacturer’s instructions. After 48 hours of transfection, the cells were used for RNA isolation and real-time PCR or reseeded on coverslips for voltage clamp measurements.

### Electrophysiology experiment

For electrophysiology experiments, the GLUTag cells were transfected with siRNA and cultured in 35mm dishes for 48 hours prior to the experiments. The experiments were performed on individual cells. The GLUTag cells were whole-cell patch-clamped using an EPC-10 amplifier and PULSE software (HEKA Electronik, Lambrecht, Germany). The Kv currents were recorded from a holding potential of -e (in mmol/L): 10 KCl, 125 NaCl, 2 CaCl2, 2 MgCl2, 1 glucose, and 10 HEPES (pH 7.4), while the intracellular solutions contained (in mmol/L): 130 KCl, 10 NaCl, 2 CaCl2, 10 EGTA, 10 HEPES, 2 MgCl2, and 1 MgATP (pH 7.4). Action potentials were elicited by a 0.5 nA current injection for 50 milliseconds in the GLUTag cells. The electrode solutions for recording action potentials contained (in mmol/L): 76 K2SO4, 10 KCl, 10 NaCl, 55 sucrose, 1 MgCl2, and 10 HEPES (pH 7.2), while the extracellular solutions contained (in mmol/L): 5.6 KCl, 138 NaCl, 2.6 CaCl2, 1.2 MgCl2, 1 glucose, and 10 HEPES (pH 7.4). The data were recorded using pCLAMP software (Axon Instruments) and analyzed using ClampFit software. The experiments were performed at a temperature of 22-24°C.

### GLP-1 secretion assay

GLUTag cells were plated in 24-well plates at a density of 200,000 cells per well in 1 ml of culture medium. After 48 hours, the cells were rinsed with HBSS and incubated in Krebs–Ringer Bicarbonate Buffer (KRBB; 120 mM NaCl, 5 mM KCl, 24 mM NaHCO^3^, 1 mM MgCl2, 2 mM CaCl2, 15 mM HEPES pH 7.4, 0.1% BSA) for 30 minutes. The buffer was then replaced with either KRBB or high-glucose KRBB containing 10 mM glucose and the cells were incubated for an additional 2 hours. Some GLUTag cells were treated with 10 mM glucose KRBB containing 10 μM Forskolin (Sigma), 10 μM IBMX (Sigma), 2 μM OEA (MCE), or 10 μM Glutamine(Gibco). Following a 2-hr incubation, the media was collected and the cells were lysed in lysis buffer. GLP-1 levels were measured using a total GLP-1 ELISA kit (Millipore) and an Infinite 200 Pro Reader (TECAN, Switzerland).

### Measurement of [Ca2+]i

Cells were plated on glass coverslips and left in a humidified incubator for 24 hours. Before imaging, the cells were loaded with 2 μM Fura-4 AM (Dojindo, Japan) in KRBB (0 mM glucose) at 37°C for 30 minutes, then washed twice with low-glucose KRBB without Fluo4-AM. The cells were then pre-incubated with 0 mM glucose KRBB for an additional 30 minutes and stimulated with 10 mM high-glucose KRBB. The experiments were performed using a DeltaVision Ultra High Resolution Microscope (GE, USA) with a 60× objective. The fluorescence was excited at 488 nm and the emission was collected at 525 nm. Sequential confocal images of the cells were recorded every 5 seconds starting 60 seconds before stimulation and analyzed using Image J software. The ratio of fluorescence change (F/F0) was used to reflect changes in intracellular Ca2+ levels, where F represents the observed fluorescence density and F0 represents the average baseline fluorescence density of the first 30 seconds before stimulation.

### Statistical analysis

Data are presented as means ± SEM values and compared using a Student’s t-test. A p-value of less than 0.05 was considered statistically significant.

## Results

### HERG was expressed in the murine intestinal L-cells

To determine its expression level, the mRNA expression of HERG was analyzed in the murine ileum, colon and GLUTag cells using RT-PCR (Figure 1A). This expression was further confirmed by western blot analysis, which showed that HERG protein was highly expressed in GLUTag cells and even more so than in murine brain, which was used as a positive control (Figure 1B). To study the expression of the Kv channels in GLUTag cells, a qPCR profile was performed (Figure 1C). The results showed that Kv2.1, -4.1, -5.1, -6.2, and -9.3, in addition to Kv11.1, were highly expressed, with Kv11.1 (HERG1) being the second highest. To determine the distribution of HERG, immunofluorescence was performed on frozen sections of murine ileum tissue. The results showed that HERG colocalized with GLP-1 on the intestinal L-cells (Figure 1D).

**Figure 1.**
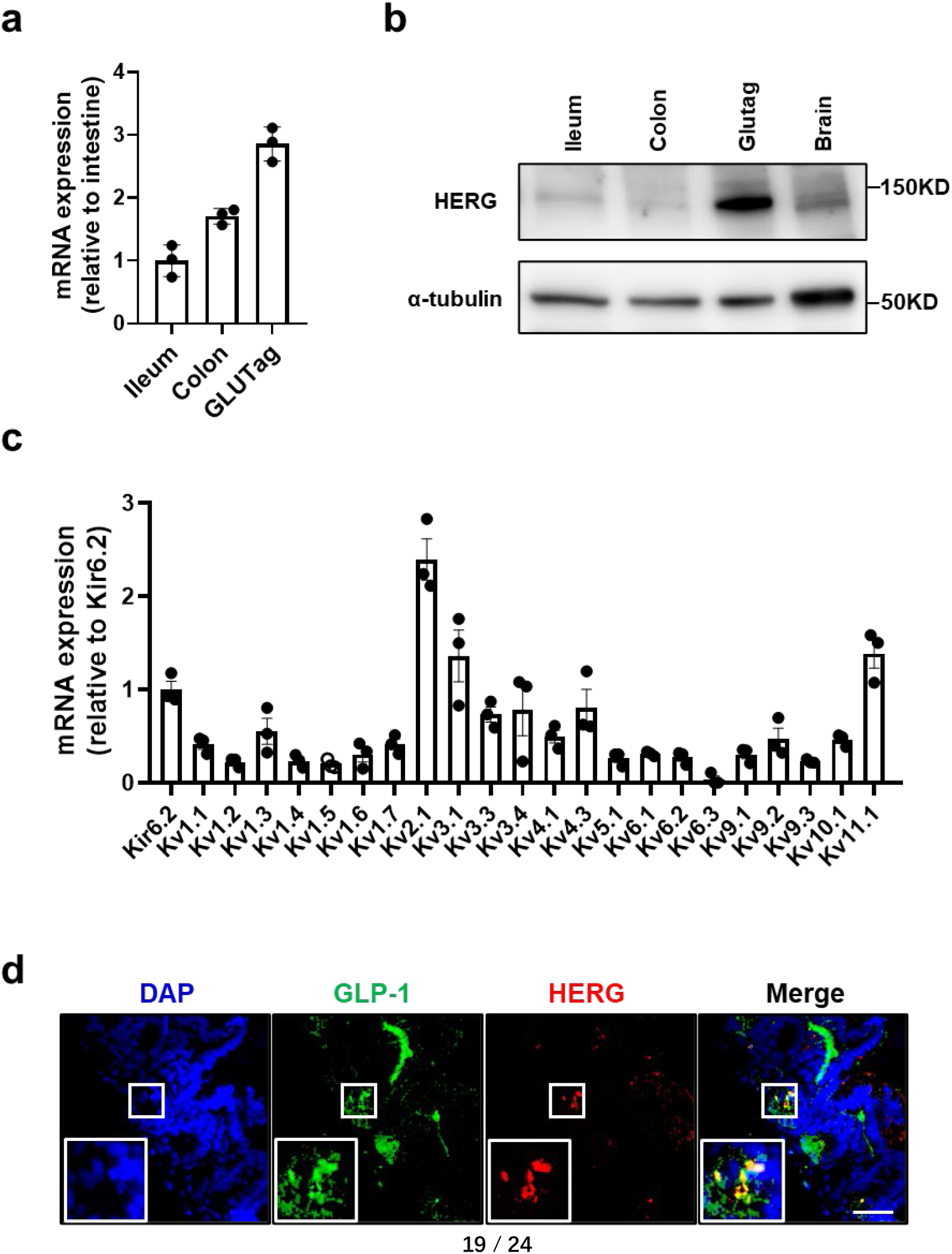
Expression of HERG in GLUTag cells and murine L-cells. (a) Quantitative PCR analysis of mRNA expression of HERG in murine ileum, colon and GLUTag cells. (b) Western blot analysis of protein expression of HERG in murine ileum, colon, brain and GLUTag cells. (c) mRNA expression of Kv channels in GLUTag cells, expressed as a percentage of Kir6.2 (KATP channels) after normalization with internal control. (d) Immunofluorescence analysis of localization of HERG channel (green) in murine ileum, with GLP-1 (red) as the marker of L-cells. Nuclei (blue) were stained with DAPI. Scale bar represents 50 μm. Each experiment was repeated at least three times.

### The effect of HERG on Kv current and action potential in GLUTag cells

HERG expression was suppressed in GLUTag cells by using siRNA. qPCR and western blot analysis confirmed that HERG expression was significantly decreased compared to the control (scramble siRNA) (Fig. 2B). To assess the impact of HERG on the whole-cell Kv currents, patch-clamp recordings were performed as described in the methods section (Fig. 2C). The analysis of e showed that HERG siRNA-treated GLUTag cells displayed a significant reduction in Kv currents compared to the control cells (Fig. 2D). The Kv channels play a crucial role in the repolarization of action potentials in pancreatic islet β-cells, and the inhibition of Kv channels results in an extended action potential duration (APD). Electrical action potentials were evoked in current-clamp mode to determine the APD in GLUTag cells, and the results showed that the repolarization of action potentials was prolonged in HERG siRNA-treated cells compared to control cells (Fig. 2E). These findings demonstrate that the HERG channel regulates the repolarized Kv current and that the reduction of the HERG channel leads to a prolonged APD in GLUTag cells.

**Figure 2.**
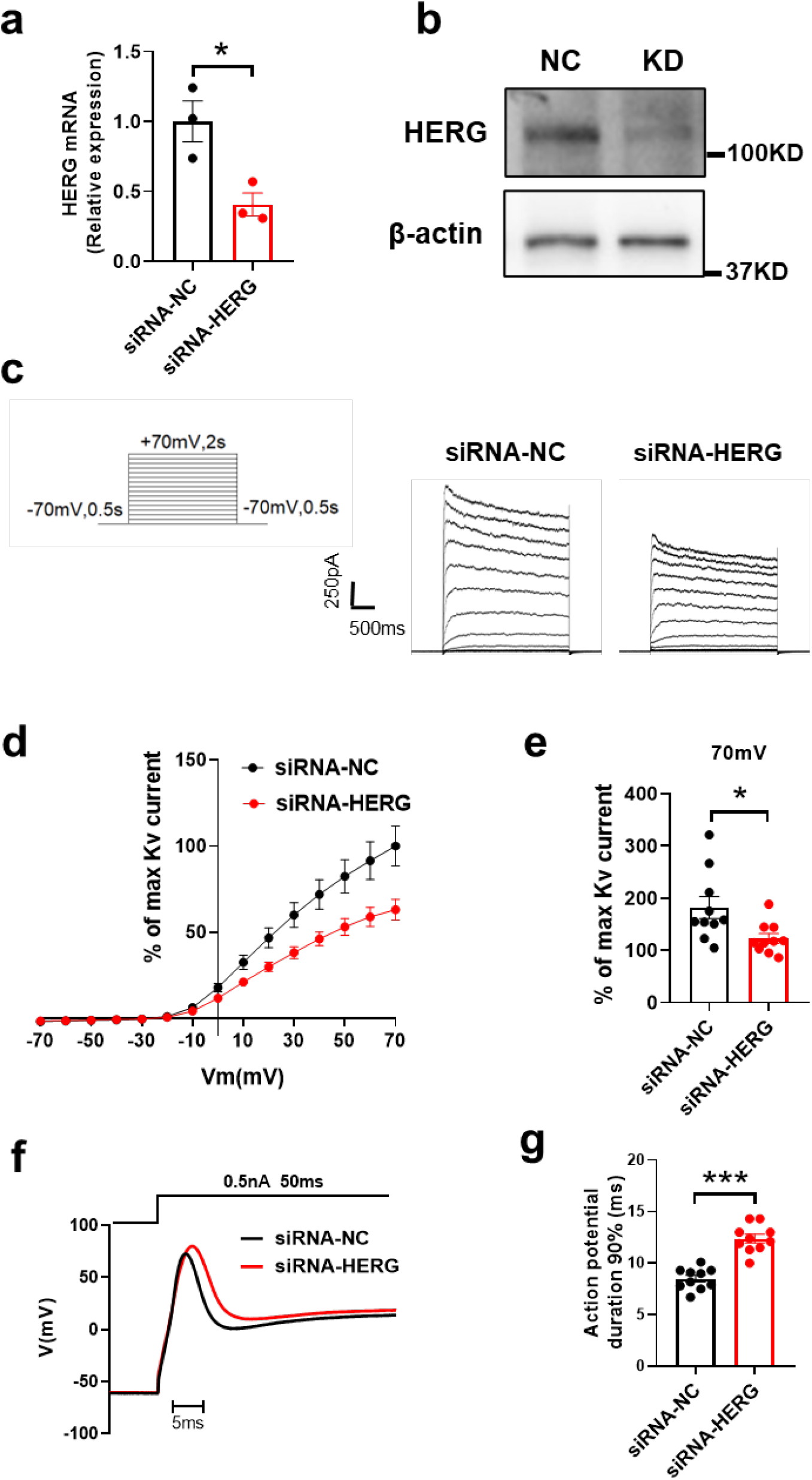
Prolongation of action potential duration in GLUTag cells upon knockdown of HERG channel. (a, b) GLUTag cells were transfected with either 100 nM scramble siRNA (siRNA-NC) or siRNA against HERG (siRNA-HERG) for 48 hours, and HERG mRNA (a) and protein (b) expression of HERG were evaluated using qPCR and western blot analysis (n = 3). (c) Representative whole-cell recordings of Kv currents from GLUTag cells treated with either siRNA-NC or siRNA-HERG. (d) The cells were depolarized in increments of 10 mV from a holding potential of -70 mV for a duration of 500 ms, and the steady-state current-voltage curves for Kv currents were determined by normalizing the data to the maximum current (siRNA-NC, n = 10; siRNA-HERG, n = 10). (e) Bar graphs displaying the Kv current at a potential of 70 mV. (f) Representative action potentials recorded from GLUTag cells in current-clamp mode. (g) Bar graphs exhibiting action potential duration (siRNA-NC, n = 10; siRNA-HERG, n = 10). Statistical significance was determined using a Student t-test, and values denoted by *p < 0.05 and ***p < 0.001 were considered statistically significant.

### The impact of HERG on calcium concentration ([Ca2+]i) in GLUTag cells

To further understand the role of HERG channels in calcium homeostasis, we examined the effect of HERG on the [Ca2+]i in GLUTag cells. Using siRNA, we reduced HERG expression in GLUTag cells and measured the [Ca2+]i response to 10 mM glucose. Our results showed that HERG reduction significantly increased the [Ca2+]i (Figure 3A) with a 1.44-fold increase compared to control cells (Figure 3B). This suggests that HERG reduction enhances [Ca2+]i after glucose stimulation in GLUTag cells.

**Figure 3.**
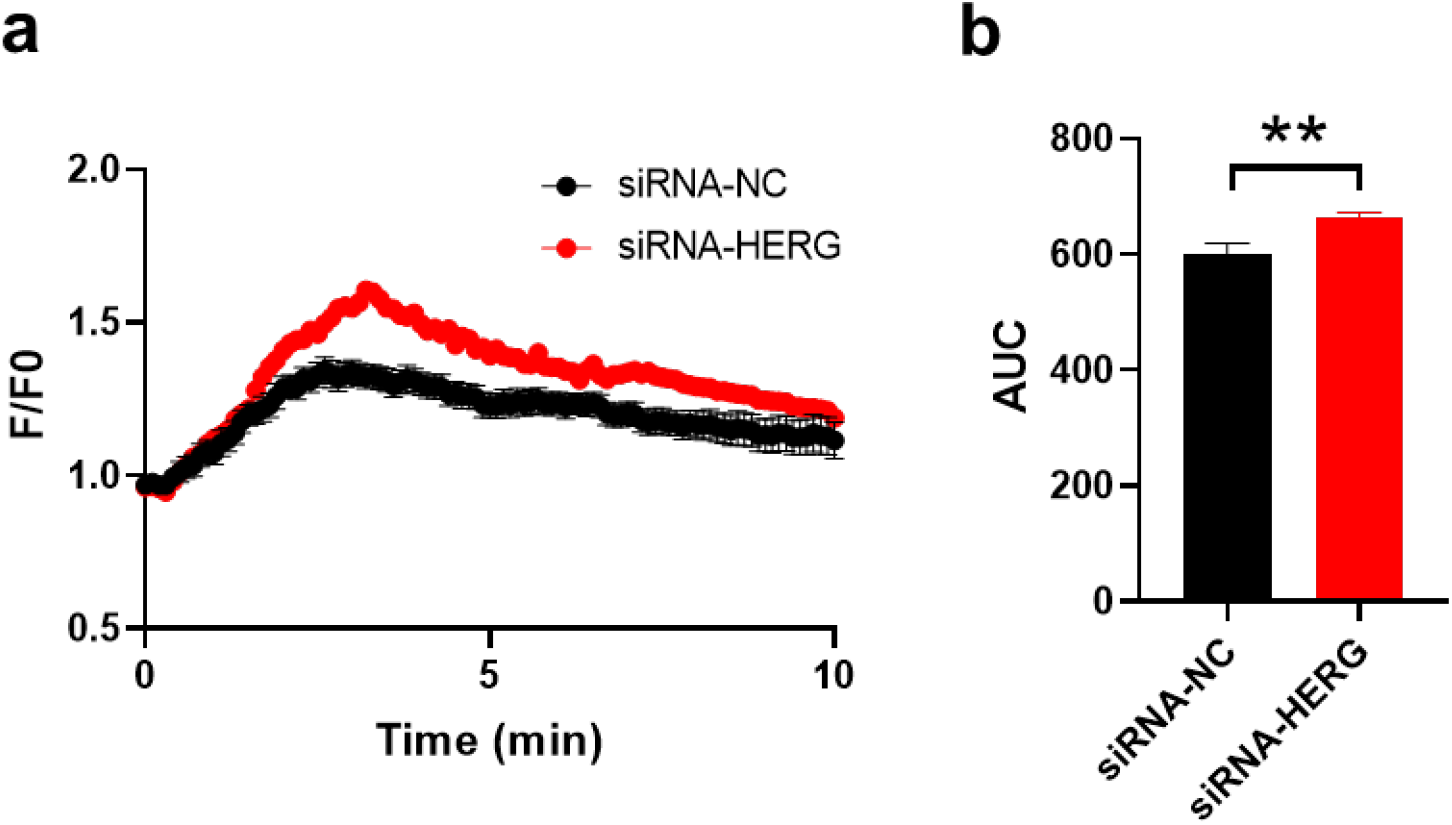
The effect of HERG knockdown on intracellular calcium concentration ([Ca2+]i) in GLUTag cells. [Ca2+]i was determined using 2 μM Fluo 4-AM and a DeltaVision Ultra High Resolution Microscope equipped with a 60× objective, at a temperature of 37°C. Readings were taken every 5 seconds for a total of 31 minutes, with 1 minute being recorded prior to and 30 minutes following glucose stimulation (10 mM). (a) The fluorescence change ratio (F/F0) from GLUTag cells treated with scramble siRNA (n=23 cells) and HERG siRNA (n=23 cells) was recorded. (b) The average area under curve (AUC) of the HERG siRNA group was found to be significantly higher compared to the control group (p<0.01). The statistical significance of the difference between means was assessed using a Student’s t-test. \*\**p*<0.01.

### The effect of HERG on GLP-1 secretion in GLUTag cells

Given that HERG reduction increased intracellular [Ca2+], we investigated whether it would also enhance GLP-1 secretion in GLUTag cells. We found that while HERG reduction had no effect on GLP-1 secretion under basal conditions (0 mM glucose), it significantly increased GLP-1 secretion after 10 mM glucose stimulation (Figure 4A). To further test the effect of other stimulators, we also stimulated GLUTag cells with 10 mM Glutamine to mimic amino acid stimulation, 2 μM GPR119 agonist 2-oleoylglycerol (OEA) to mimic fatty acid stimulation, and 10 μM forskolin plus 10 μM 3-isobutyl-1-methylxanthine (IBMX) to mimic cAMP stimulation. Our results showed that Glutamine and cAMP stimulation significantly enhanced GLP-1 secretion in HERG reduction cells (Figure 4B, D), while OEA stimulation showed only a slight increase in GLP-1 secretion (Figure 4C). These findings suggest that HERG reduction enhances GLP-1 secretion in GLUTag cells.

**Figure 4.**
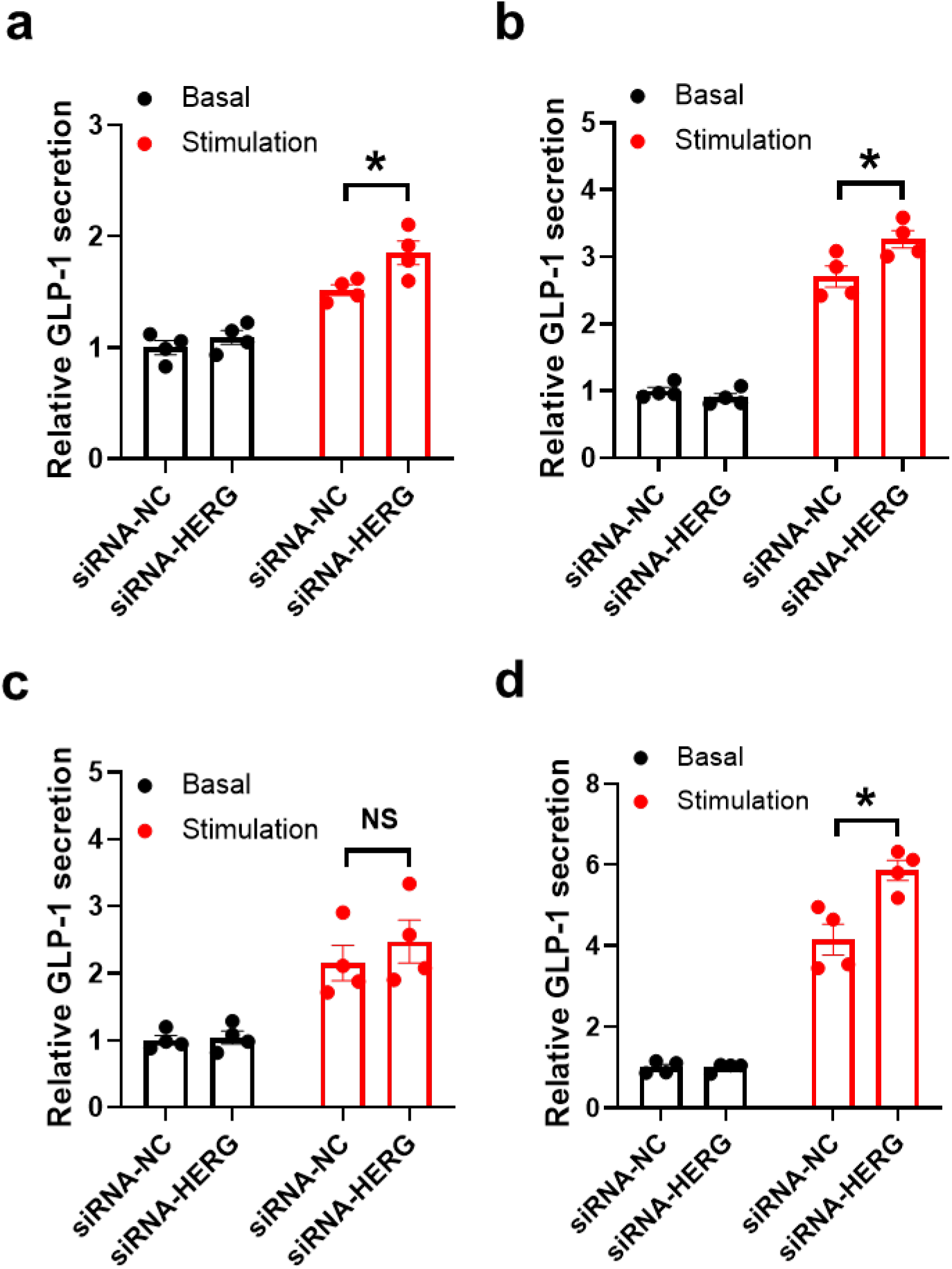
HERG knockdown enhances GLP-1 secretion in GLUTag cells. The GLUTag cells were treated with 10 mM glucose (a), 10 mM glutamine (b), 2 μM OEA (c), and 10 μM forskolin plus 10 μM IBMX (d) individually. The results demonstrate that the reduction of HERG expression significantly stimulates GLP-1 secretion from GLUTag cells. Statistical significance was determined using Student’s t-test, with *p<0.05 indicating a significant difference between the means.

## Discussion

In this study, we demonstrated the expression of HERG potassium channels in the mice ileum, colon, and the e GLUTag cells. Upon downregulation of HERG, we observed a prolonged action potential duration and an increase in intracellular calcium concentration. Furthermore, our findings revealed that the deficiency of HERG promoted the secretion of GLP-1 in response to glucose and amino acid stimulation. These results suggest the involvement of HERG in the release of GLP-1 from the intestinal L-cells.

It has been reported that the nutrient-stimulated secretory pathways in L-cells and β cells share a common glucose sensing mechanism. The intake of nutrients such as glucose and amino acids leads to an increase in ATP production and a subsequent closure of KATP channels, resulting in the depolarization of the plasma membrane (12, 20). This leads to the influx of calcium, which triggers hormone exocytosis during action potential firing (12, 20). Kv channels, which play a crucial role in regulating hormone secretion (11), also open during depolarization, thus repolarizing the action potential. GLP-1 is secreted by intestinal L-cells in response to dietary components, including carbohydrates, amino acids, and fats (21, 22). These cells are distributed throughout the gut, but are most abundant in the ileum and colon (23, 24). With their apical protrusions facing the gut lumen, L-cells are believed to directly sense the concentration of fats and carbohydrates in the lumen. The glucose sensing mechanism in L-cells is similar to that in pancreatic β cells. As such, we conducted a quantitative polymerase chain reaction screening for Kv channel isoforms and found that the HERG channels were the second-highest expressed in the murine L-cell line. This finding suggests that the HERG potassium channel may play a role in regulating the secretion of GLP-1 from L-cells.

Mutations in HERG can reduce potassium efflux during repolarization, resulting in prolonged QT interval and long QT syndrome (LQTS) (25). In LQT2 patients, HERG mutations have been reported to increase GLP-1 and insulin secretion by over 50%, as well as causing defects in glucagon secretion. This can lead to an increased risk of low blood sugar levels and symptomatic hypoglycemia following glucose ingestion. The HERG receptor blocker, dofetilide, has a similar effect by inhibiting HERG in both β and L, leading to increased insulin and GLP-1 secretion (2).

L-cells are open enteroendocrine cells with apical protrusions facing the intestinal lumen, believed to play a role in nutrient sensing. They respond to a range of luminal components, particularly the products of carbohydrate, fat and protein digestion (21, 22). It is proposed that electrical uptake of glucose or amino acids triggers membrane depolarization, electrical activity, calcium influx via voltage-gated calcium channels, and GLP-1 secretion in L-cells (17, 26-28).

Our study of patch-clamp electrophysiology combined with HERG gene silencing in GLUTag cells revealed a reduction in HERG channels, which prolonged action potential durations, inducing more calcium influx. Calcium imaging also showed increases in intracellular calcium [Ca2+]i after glucose stimulation in HERG downregulated GLUTag cells. We also found that downregulation of HERG enhanced GLP-1 secretion in response to nutrient stimulation in GLUTag cells. These findings support the role of HERG channels in repolarizing intestinal L-cells and demonstrate the critical role of HERG potassium channel activity in regulating GLP-1 secretion. Further studies using HERG L-cell specific knockout mice will clarify the in vivo function of HERG in GLP-1 secretion.

## Declarations

### Ethics approval and consent to participate

All animal protocols were approved by the Ethical Review Committee at the Institute of Zoology, Capital Medical University, China.

### Consent for publication

Not applicable.

### Availability of data and material

JKY is the guarantor of this work and, as such, the data that support the findings of this study are available from him upon reasonable request.

## Authors’ contributions

JKY conceived the idea for the study. CL, YCY, RRX and HW performed the experiments and processed statistical data. CL, HW drafted the manuscript. JKY and HW is responsible for supported the study, designed the experiments, and summarizing all research data.

